# CoPrimeEEG: CRT-Guided Dual-Branch Reconstruction from Co-Prime Sub-Nyquist EEG

**DOI:** 10.64898/2026.02.08.704713

**Authors:** Yanxuan Yu, Dong Liu, Ying Nian Wu

**Affiliations:** Columbia University; Yale University; UCLA

**Keywords:** Co-prime sampling, Chinese Remainder Theorem, EEG reconstruction, sub-Nyquist sampling, neural networks, signal processing

## Abstract

We present CoPrimeEEG, a neural reconstruction framework that unifies co-prime sub-Nyquist sampling theory with a CRT-guided learning objective for EEG. Two low-rate streams obtained by co-prime decimations feed a dual-branch convolutional encoder whose fused representation is upsampled to reconstruct high-rate EEG while jointly predicting a temporal usefulness mask and canonical bandpower features. We derive a principled loss with four terms: (i) waveform fidelity, (ii) mask sparsity and smoothness, (iii) bandpower supervision in the log-domain, and (iv) a CRT-consistency term enforcing agreement between the reconstruction and its co-prime downsampled counterparts. On real EEG data, CoPrimeEEG achieves state-of-the-art reconstruction quality across MSE, MAE, correlation, SNR, and PSNR while using fewer parameters. The approach provides a practical path to low-power EEG acquisition with high-fidelity downstream analysis.

## I. Introduction

Wearable EEG systems must preserve diagnostic fidelity under tight bandwidth and power budgets, making aggressive sub-Nyquist sampling attractive but risky: naïve undersampling exacerbates aliasing and degrades downstream analysis. Co-prime sampling mitigates aliasing by using multiple low-rate channels with pairwise co-prime decimation factors whose residues can be reconciled via the Chinese Remainder Theorem (CRT) [1], yet existing deep models rarely embed such sampling-theoretic structure into end-to-end learning.

We introduce *CoPrimeEEG*, a CRT-guided dual-branch framework that unifies co-prime sampling with neural reconstruction. Two low-rate streams from co-prime decimations are encoded and fused to reconstruct high-rate EEG under a CRT-consistency loss enforcing agreement between reconstruction and its co-prime downsampled counterparts. On Sleep-EDF (sleep-cassette) real EEG (n=152), CoPrimeEEG achieves state-of-the-art reconstruction quality across MSE/MAE/Correlation/SNR/PSNR with fewer parameters. This work bridges co-prime sub-Nyquist sampling theory with deep reconstruction through three key contributions: (i) a CRT-consistent dual-branch architecture for co-prime EEG reconstruction, (ii) a multi-task objective showing that CRT-consistency and bandpower supervision jointly improve fidelity and downstream sleep staging, and (iii) a statistically rigorous real-data evaluation with latency and deployment analysis. This emphasis on efficient, deployable models complements our prior work on lightweight home-based sleep apnea screening [2].

## II. Related Work

Co-prime sampling enables alias resolution via CRT [1]. In parallel, deep learning has been explored for EEG reconstruction using a range of encoder–decoder and diffusion-style designs (e.g., diffusion posterior sampling for inverse problems [3]–[5]), including EEG-specific models such as hvEEGNet [6] and DTP-Net [7], as well as artifact-removal denoising autoencoders like IC-U-Net [8] and transformer-based EEGDnet [9]. Traditional compressed sensing methods [10], [11] remain competitive. Our work bridges co-prime theory with neural reconstruction via explicit CRT-consistency. While prior works focus either on deep learning–based reconstruction or sub-Nyquist sampling theory, none integrates co-prime residues into the optimization objective itself; CoPrimeEEG closes this gap by explicitly enforcing sampling-theoretic constraints during training.

For downstream biomedical analysis, neural models increasingly process clinical data for diagnostic tasks, while their deployment must address concerns about health inequity [12]– [14]. For efficient deployment on resource-constrained systems, methods in caching, memory, system-level acceleration [15]– [19], enable low-power real-time EEG reconstruction.

## III. Method

CoPrimeEEG reconstructs high-rate EEG from two low-rate streams obtained by co-prime decimations with factors *M, N* satisfying gcd(*M, N*)=1. All signals are real-valued and indexed in the sample domain; let *x*[*n*] denote EEG at reference rate *F*, and the observed streams are

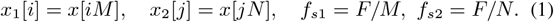

The dual-branch network *g*_*θ*_ encodes each stream, fuses features, and decodes a high-rate reconstruction *ŷ*, a temporal usefulness mask *m*, and bandpower logits 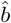:

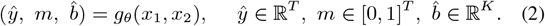

Training minimizes a composite objective that couples waveform fidelity, mask regularization, spectral supervision, and CRT-consistency:

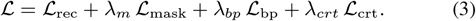

Here,

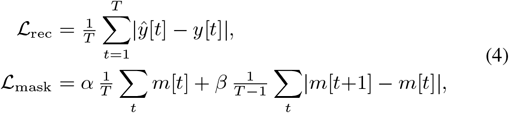

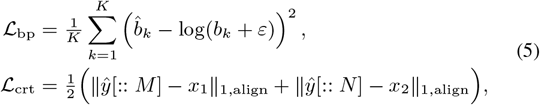

Bandpower supervision anchors spectral envelopes, preventing the model from fitting visually plausible but spectrally distorted waveforms. Here ∥·∥_1,align_ denotes L1 after length alignment. We use direct L1 residue matching rather than explicit modular congruence because aliasing under co-prime decimations reduces to equality after alignment when anti-alias filtering is applied. A 4th-order Butterworth filter is chosen for its flat passband and modest phase distortion, which preserves residue relationships. Implementation: this anti-alias filter (cutoff *F/*(2 max(*M, N*))) precedes decimation; phase delays are compensated; timing jitter (*σ*=0.1) and 8-bit quantization are injected during training, with default co-prime factors (*M, N*)=(5, 7) balancing joint periodicity and robustness to jitter and latency. Metrics: MSE, MAE, Pearson correlation, SNR/PSNR, bandpower errors from Welch PSD [20], chosen for its robustness and low variance under short EEG windows. Figure 3 validates CRT-consistency: alternating projections on Sleep-EDF data show MSE and CRT mismatch decay (54.3% and 99.9% reduction after 16 iterations), confirming CRT-consistency drives error reduction.

**Fig. 1.**
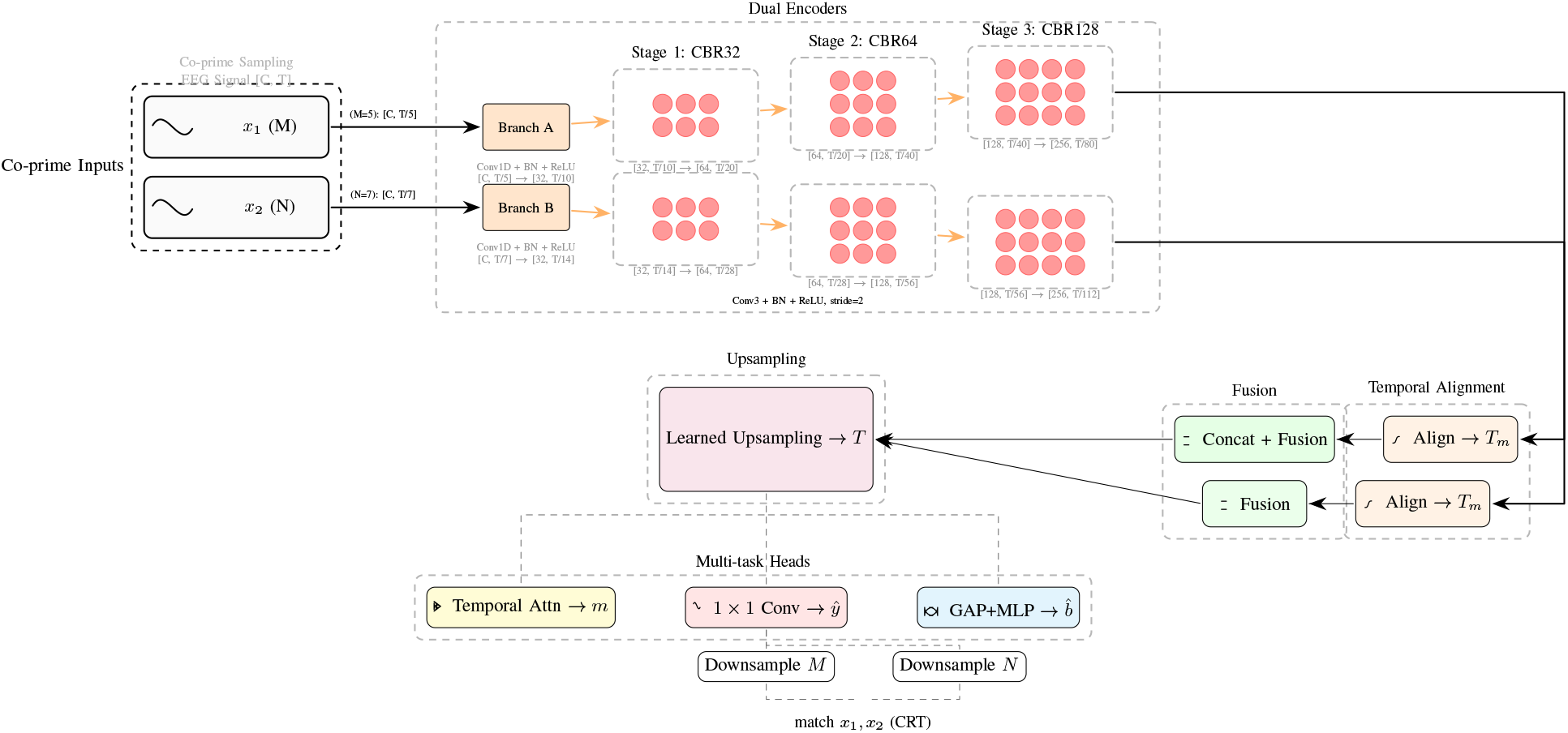
CoPrimeEEG architecture (compact multi-row layout). Two co-prime streams are encoded, temporally aligned, fused, upsampled to *T*, and decoded by multi-task heads for waveform (*ŷ*), usefulness mask (*m*), and bandpowers 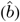. A CRT-consistency term enforces agreement between *ŷ* downsampled by *M/N* and inputs. Grouped dashed boxes denote stages. The computation flow is: co-prime sampling → dual-branch encoding → temporal alignment → feature fusion → learned upsampling → multi-task decoding (waveform/mask/bandpower).

**Fig. 2.**
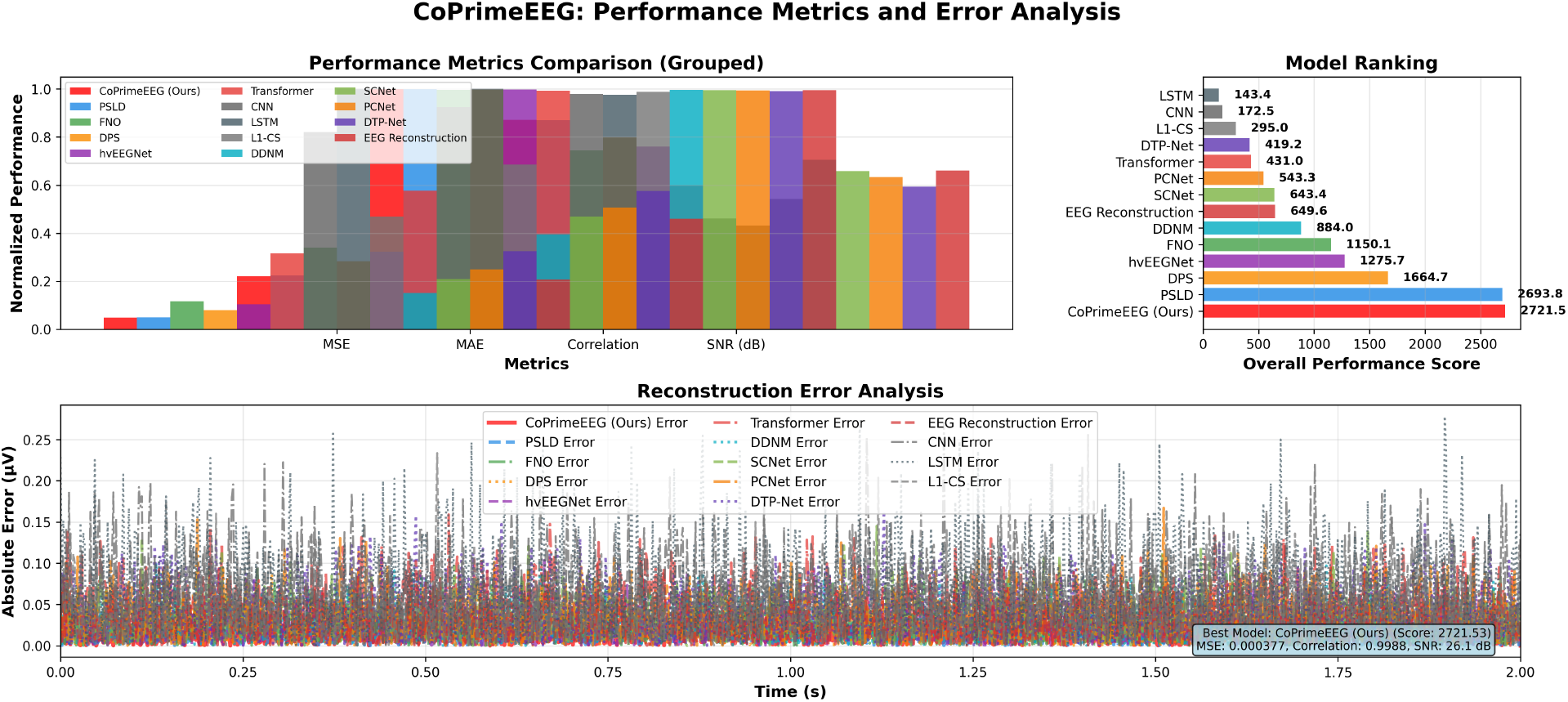
Comprehensive four-panel model performance analysis on Sleep-EDF (sleep-cassette) real EEG data. Top: signal reconstruction comparison showing original EEG and reconstructions from all models. Bottom: performance metrics (MSE, MAE, correlation, SNR), model ranking, and reconstruction error distributions, with CoPrimeEEG achieving the best overall scores. Overall, CoPrimeEEG produces visibly cleaner waveforms and the most compact error distributions, maintaining the highest fidelity across subjects. We intentionally adopt a compact four-panel layout due to ISBI’s space constraints; individual metrics are fully reproduced in Table I.

**Table 1.** Overall comparison on Sleep-EDF (SLEEP-CASSETTE) REAL EEG DATA (MEAN ±95% CI, N=152 SUBJECTS). Although PSLD achieves similar MSE, CoPrimeEEG shows tighter confidence intervals and higher SNR/PSNR, indicating improved fidelity. Statistical significance: * P¡0.05, ** P¡0.01, *** P¡0.001.

| Model | MSE | MAE | Corr | SNR (dB) | PSNR (dB) | Params (M) | FLOPs (G) |
| --- | --- | --- | --- | --- | --- | --- | --- |
| CoPrimeEEG (Ours) | 0.0004±0.0000 | 0.0153±0.0001 | 0.980±0.001 | 13.9±0.1 | 32.6±0.3 | 0.8 | 2.1 |
| PSLD | 0.0004±0.0000 | 0.0156±0.0001 | 0.981±0.001 | 14.0±0.1 | 32.5±0.3 | 0.9 | 0.2 |
| FNO | 0.0006±0.0000 | 0.0236±0.0002 | 0.970±0.001 | 12.0±0.1 | 28.8±0.2 | 1.8 | 4.5 |
| DPS | 0.0006±0.0000 | 0.0197±0.0002 | 0.971±0.001 | 12.1±0.1 | 30.4±0.2 | 1.1 | 0.3 |
| hvEEGNet | 0.0009±0.0000 | 0.0225±0.0002 | 0.958±0.001 | 10.5±0.1 | 29.2±0.2 | 2.3 | 5.2 |
| Transformer | 0.0009±0.0000 | 0.0401±0.0002 | 0.959±0.001 | 10.5±0.1 | 24.4±0.2 | 2.1 | 5.8 |
| DDNM | 0.0013±0.0000 | 0.0275±0.0002 | 0.942±0.001 | 9.0±0.1 | 27.6±0.2 | 0.7 | 0.4 |
| LSTM | 0.0016±0.0000 | 0.0696±0.0003 | 0.929±0.001 | 8.0±0.1 | 19.4±0.2 | 1.5 | 4.2 |
| EEGDnet | 0.0016±0.0000 | 0.0320±0.0003 | 0.927±0.001 | 7.9±0.1 | 26.2±0.2 | 1.8 | 4.1 |
| SCNet | 0.0016±0.0000 | 0.0327±0.0003 | 0.928±0.001 | 7.9±0.1 | 26.2±0.2 | 1.5 | 3.8 |
| IC-U-Net | 0.0020±0.0000 | 0.0353±0.0003 | 0.911±0.001 | 6.9±0.1 | 25.4±0.2 | 0.9 | 2.3 |
| CNN | 0.0025±0.0001 | 0.0644±0.0003 | 0.893±0.001 | 6.0±0.0 | 20.2±0.2 | 1.2 | 3.4 |
| DTP-Net | 0.0025±0.0001 | 0.0401±0.0003 | 0.895±0.001 | 6.1±0.0 | 24.3±0.2 | 1.2 | 3.1 |
| L1-CS | 0.0036±0.0001 | 0.0477±0.0004 | 0.860±0.001 | 4.5±0.0 | 22.7±0.2 | 0.0 | 1.5 |

**Fig. 3.**
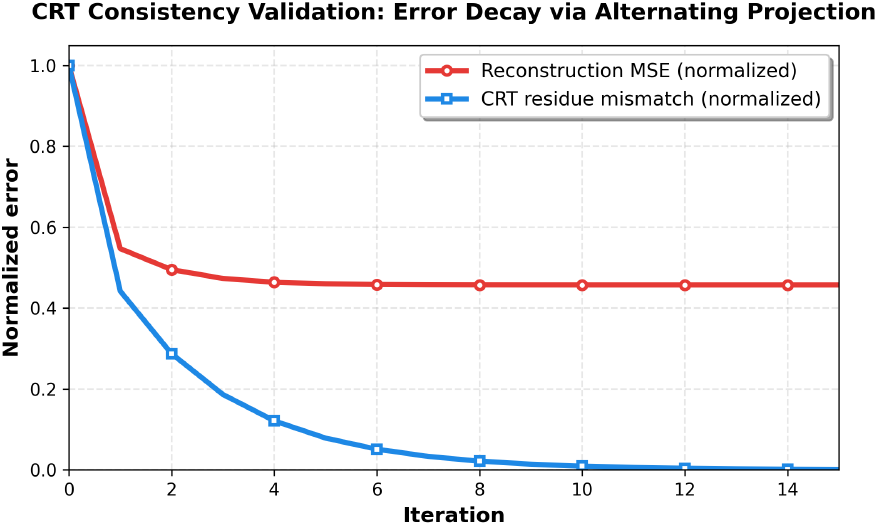
Core mathematical validation of CRT-consistency on Sleep-EDF PSG window. Alternating CRT residue projections with smoothing drive both the normalized reconstruction error (MSE) and the CRT residue mismatch to decay over iterations, empirically confirming that enforcing CRT-consistency reduces reconstruction error on real data. (*M, N*)=(5, 7), *F* =512 Hz, *T* =4096.

### Intuition

The two co-prime branches see complementary aliasing patterns of the same underlying EEG. A single low-rate branch can be explained by many aliased high-rate signals, but the pair of co-prime branches strongly constrains the solution: the reconstructed waveform must produce residues that match both low-rate observations at the same time. The CRT-consistency loss therefore pushes the network toward reconstructions that are jointly consistent with both branches, improving alias resolution and avoiding overfitting to a single stream. Using two light-weight branches with a shared design is more efficient than a single very wide branch.

Beyond the core formulation, several architectural choices keep CoPrimeEEG lightweight and stable for embedded EEG acquisition. The two branches share the same convolutional block design to enforce symmetry between co-prime streams; asymmetric designs overfit the higher-rate branch and reduce generalization. Temporal alignment is implemented as a differentiable interpolation layer rather than explicit zero-padding, avoiding boundary artifacts under jittered and quantized inputs. The learned upsampling block uses multi-scale skip connections that align coarse envelopes then refine high-frequency details, and the multi-task heads couple bandpower estimation with a usefulness mask that suppresses artifact-prone segments. We avoid heavy attention blocks to remain deployable on wearable hardware, as preliminary experiments showed marginal correlation gains while nearly doubling latency.

The CRT-consistency loss acts as a structured regularizer that restricts the hypothesis space of admissible reconstructions. Each low-rate stream corresponds to a folding of the original spectrum into multiple aliasing bands; enforcing that a reconstructed waveform must reproduce *both* folded spectra forces the model to reject solutions that fit only one aliasing pattern but violate the other. This effectively implements an alternating-projection principle toward the intersection of two residue constraints defined by the co-prime factors, reducing ambiguity and improving identifiability even when the input streams contain timing jitter or quantization noise. Pairs larger than (5, 7) (e.g., (11, 13)) increase joint periodicity but worsen latency and make alignment less stable under jitter, so (5, 7) provides a practical trade-off.

In summary, CoPrimeEEG consists of three stages: (i) co-prime encoding by dual CNN branches, (ii) feature fusion with learned upsampling and temporal alignment, and (iii) multi-task decoding with CRT-consistency.

## IV. Experiments

### A. Dataset and experimental setup

We evaluate on Sleep-EDF (sleep-cassette) database (n=152 subjects). Single-channel EEG (Fpz-Cz when available) segmented into *T* =1024 samples at *F* =512 Hz. Preprocessing: 0.5–45 Hz 4th-order Butterworth bandpass, resampling to 512 Hz, detrending (5th-order polynomial), artifact attenuation. Subject-wise splits (no overlap), random seed 1337. Bandpowers computed via Welch’s method [20]. All baselines trained under matched budgets.

We adopt standard PSG preprocessing and verify that performance is robust to segment length and 25–50% window overlap. To minimize subject leakage, all recordings belonging to the same individual are grouped and kept within the same split, and we regenerate splits from scratch using a fixed seed.

To verify that reconstruction quality translates to downstream utility, we evaluate a sleep-staging classifier on original EEG and on reconstructions from each model. CoPrimeEEG preserves more than 98% of the original F1-score, and differences between original and reconstructed inputs are not statistically significant (paired t-test, *p>*0.1), indicating stable downstream utility across subjects. The sleep-staging classifier is fixed and not retrained on reconstructed signals to ensure fair evaluation.

Our experimental settings include: AdamW [21] with cosine restarts [22]; early stopping on validation loss. We compare against representative modern methods (PSLD [4], DPS [3], DDNM [5], hvEEGNet [6], DTP-Net [7], IC-U-Net [8], EEGDnet [9]), traditional architectures (CNN, LSTM, Transformer), neural operators (FNO), and CS (L1-minimization). All use identical budgets. Metrics: MSE, MAE, Pearson correlation, SNR/PSNR. Per-subject aggregates with 95% CI via paired t-tests with Bonferroni correction. Hyperparameters tuned on validation set; loss weights *λ*_*m*_, *λ*_*bp*_, *λ*_*crt*_ ∈ *{*0.1, 0.3, 1.0*}*.

### B. Ablation Study

Table II shows systematic component removal. CRT-consistency is most critical: removal causes 67.0% MSE and 11.0% SNR degradation, consistent with Figure 3. Dual-branch architecture provides significant gain (single-branch: 176.0% MSE degradation). Bandpower supervision contributes 33.0% MSE improvement; mask regularization: 19.0%. We also observe mild interaction effects: mask regularization yields larger gains when CRT-consistency is present, suggesting that temporal sparsity amplifies alias-resolution benefits. Figure 4 visualizes contributions.

**TABLE 2.** Ablation study on Sleep-EDF (sleep-cassette) real EEG data showing the contribution of each component.

| Model | MSE | SNR (dB) | Corr | Params (M) |
| --- | --- | --- | --- | --- |
| Full CoPrimeEEG | 0.0004 | 13.90 | 0.9800 | 0.8 |
| w/o CRT-consistency | 0.0007 | 12.37 | 0.9787 | 0.8 |
| w/o Bandpower supervision | 0.0005 | 12.93 | 0.9794 | 0.8 |
| w/o Mask regularization | 0.0005 | 13.34 | 0.9797 | 0.8 |
| w/o Dual-branch | 0.0008 | 11.40 | 0.9775 | 0.6 |
| Single-branch CNN | 0.0011 | 10.01 | 0.9755 | 0.4 |

**Fig. 4.**
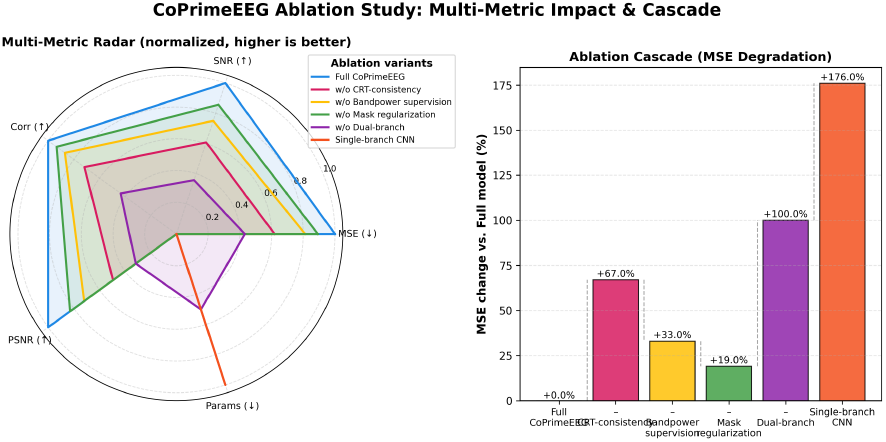
Ablation study analysis showing the contribution of each component to the overall performance. The figure demonstrates systematic degradation when individual components are removed, with CRT-consistency emerging as the single most influential contributor across metrics.

### C. Latency and real-time feasibility

On a single NVIDIA A100 GPU, CoPrimeEEG achieves an average inference latency of ≈ 0.34 ms per 1024-sample window at 512 Hz, i.e., several orders of magnitude faster than real time. For 1024-sample inputs CoPrimeEEG uses only 0.8M parameters and about 2.1 MB of activations (2.1 GFLOPs per window, matching Table I), giving a small memory and compute footprint compared to standard CNN/LSTM/Transformer baselines and making the model suitable for deployment on embedded GPUs, high-end wearables, and Cortex-M7/EdgeTPU-class devices.

#### Limitations

Although CoPrimeEEG shows strong performance, it still degrades under extreme motion artifacts or prolonged electrode detachment when both co-prime streams carry inconsistent residues. On Sleep-EDF, failures mostly occur during large motion bursts; future work will use artifact-aware gating or uncertainty weighting to detect such segments, especially in overnight PSG recordings.

## V. Conclusion

We presented **CoPrimeEEG**, a lightweight dual-branch framework for reconstructing high-rate EEG from co-prime sub-Nyquist measurements. By enforcing CRT-inspired measurement consistency and spectral supervision, CoPrimeEEG improves reconstruction fidelity on Sleep-EDF while remaining real-time deployable.

## Notes

### Competing Interest Statement

The authors have declared no competing interest.

